# RIPK3 facilitates host and pathogen interactions after oral *Toxoplasma gondii* infection

**DOI:** 10.1101/2020.08.04.237362

**Authors:** Patrick W. Cervantes, Laura J. Knoll

## Abstract

*Toxoplasma gondii* infection activates pattern recognition receptor (PRR) pathways that drive innate inflammatory responses to control infection. Necroptosis is a pro-inflammatory cell death pathway apart of the innate immune response that has evolved to control pathogenic infection. In this study we further defined the role of Z-DNA binding protein 1 (ZBP1) as a PRR and assessed its contribution to necroptosis as a host protection mechanism to *T. gondii* infection. We found that ZBP1 does not induce pro-inflammatory necroptosis cell death and ZBP1 null mice have reduced survival after oral *T. gondii* infection. In contrast, mice deleted in receptor-interacting serine/threonine-protein kinase 3 (RIPK3^-/-^), a central mediator of necroptosis, have significantly improved survival after oral *T. gondii* infection even with higher parasite burden. The physiological consequences of RIPK3 activity did not show any differences in intestine villi immunopathology but RIPK3^-/-^ mice showed higher immune cell infiltration and edema in the lamina propria. The contribution of necroptosis to host survival was clarified with mixed lineage kinase domain like pseudokinase null (MLKL^-/-^) mice. We found MLKL^-/-^ mice to succumb to oral *T. gondii* infection the same as wild type mice, indicating necroptosis-independent RIPK3 activity impacts host survival. These results provide new insights on the impacts of pro-inflammatory cell death pathways as a mechanism of host defense to oral *T. gondii* infection.

## Introduction

*Toxoplasma gondii* is one of the most widespread parasitic infections in the world and is acquired by nearly 30% of the human population (1). The natural route of infection occurs by consuming food or water contaminated with tissue cysts from the asexual cycle or oocysts from the sexual cycle. Asexual cysts contain the bradyzoite stage, which differentiates into the tachyzoite stage after ingestion and invasion of the intestinal tract. Tachyzoites disseminate from the intestine throughout the body during acute infection. Host immune pressure causes tachyzoite parasites to differentiate into bradyzoite cysts that reside in the brain and muscle tissue. This stage constitutes a latent, chronic infection. Pathogen recognition and a balanced inflammatory immune response are necessary to survive an acute infection.

*T. gondii* elicits a strong T helper type 1 (Th1) response, characterized by interleukin-12 (IL-12) induced interferon-gamma (IFN-γ), which in turn orchestrates cell-mediated immunity important for intracellular-pathogen defense (2). Deficiencies in this pathway lead to uncontrolled parasite replication and host death (1, 3, 4), highlighted in mouse studies where IL-12, IFN-γ, toll-like receptors (TLR), and myeloid differentiation primary response 88 (MyD88) responses are perturbed (5–10). These signaling pathways, cytokines, and their downstream effectors are elevated throughout long-term chronic infection and must be maintained to prevent cyst reactivation and fatal encephalitis (11, 12).

In contrast to a weak Th1 response, a hyperactive and unregulated Th1 response results in severe tissue damage and host death (13, 14). Immune regulation by IL-10 is vital to host survival because IL-10 null mice succumb to oral *T. gondii* infection due to uncontrolled host immune response and not uncontrolled parasite replication (13, 14). Early studies in susceptible C57BL/6J mice used high dose oral *T. gondii* infections and showed mortality is caused by an excessive immune response that is accompanied by severe intestinal immunopathology and tissue necrosis (15–17). This fatal pathology is mediated by the uncontrolled expression of IFN-γ, tumor necrosis factor (TNF), and nitric oxide (15, 16). Therefore, a balanced innate immune response is essential for a host to survive acute *T. gondii* infection.

The innate immune system employs pattern recognition receptors (PRRs) to identify pathogens. Common PRRs are TLRs, nucleotide oligomerization domain-like receptors (NLR), and DNA/RNA sensors like Z-DNA binding protein-1 (ZBP1). These PRRs induce programmed cell death pathways as a mechanism for host defense to infectious diseases (18, 19). Programmed cell death pathways include apoptosis, necroptosis, and pyroptosis. Apoptosis is a cell death pathway regarded as non-inflammatory, whereas necroptosis and pyroptosis are lytic forms of cell death that release highly pro-inflammatory damage associated molecular compounds (20, 21). Central to necroptosis is receptor-interacting serine/threonine-protein kinase 3 (RIPK3) and its downstream substrate mixed lineage kinase domain like pseudokinase (MLKL). Similarly, pyroptosis depends on caspase-1 activation that is mediated by NLR inflammasome components. The processes of necroptosis and pyroptosis ultimately induce cell membrane pores and lytic cell death that contribute to inflammatory immune responses, adaptive immunity, and host defense to pathogenic infection.

Our lab previously found *Zbp1* transcripts are highly expressed in the brains of mice chronically infected with *T. gondii* (22, 23). We also found ZBP1 to assist in host control of *T. gondii* infection *in vitro* and *in vivo* (24). Recently, IFN-γ has been shown to strongly induce ZBP1 to complex with RIPK3 to mediate inflammation and necroptosis as a host defense mechanism (25–28). Here, we tested the hypothesis that ZBP1-dependent necroptosis is protective for *T. gondii* infection.

## Materials and Methods

### Ethics statement

All animal use was approved by and in agreement with the Institutional Animal Care and Use Committee (IACUC) at the University of Wisconsin-Madison (UW-Madison) (protocol #M005217). Cats were treated in compliance with the guidelines set by the IACUC of the United States Department of Agriculture, Beltsville Area (protocol #15-017). Both institutions adhere to the regulations and guidelines set by the National Research Council. All mice were monitored daily for clinical signs of disease and were euthanized if symptoms were severe.

### Mouse experiments

All mice used were from the C57BL/6 background. Wild type (WT) were originally purchased from JAX but have been breed in the UW-Madison vivarium with all the other strains used for these studies. RIPK3 null (RIPK3^-/-^) mice were provided from Genentech (29). MLKL null (MLKL^-/-^) mice were provided by Dr. Doug Green at St. Jude Children’s Research Hospital (30). The RIPK3^-/-^ stain was genotyped by PCR with three primers: 5’-AGAAGATGCAGCAGCCTCAGCT, 5’-ACGGACCCAGGCTGACTTATCTC, and 5’-GGCACGTGCACAGGAAATAGC. The MLKL^-/-^ strain was also genotyped by PCR with primers: 5’-TATGACCATGGCAACTCACG, 5’-ACCATCTCCCCAAACTGTGA, and 5’-TCCTTCCAGCACCTCGTAAT.

### Z-DNA Binding Protein-1 CRISPR-Cas9 generated mouse knockout

The ZBP1 null (ZBP1^-/-^) mouse was generated at the UW-Madison Biotechnology Center. Two guide RNAs were designed to target the 5’-end (CGATCCCCTCTTACGTAATA; Chr:2 173219317-173219336) and 3’-end (TCAATCAATCGATCAACCGC; Chr:2 173207132-173207151) of *Zbp1*, removing 14KB of genomic DNA (gDNA) that included the promoter and all splice variants (Fig. S1A). The RNA guides were optimized for the least off-target effects within the Zhang Laboratory CRISPR Design tool (crispr.mit.edu). The guides were cloned into the plasmids pX330 or pX548 (obtained from Zhang Laboratory via Addgene; Cambridge, MA) and the PCR products were transcribed with the T7 MEGAshortscript kit (Life Technologies). A mixture of 50 ng/μL guide RNAs and 40 ng/μL of Cas9 protein in injection buffer (5 mM Trizma Base, 5 mM Tris HCL, 0.1 mM EDTA, pH 7.4) was injected into the pronucleus of one-day old fertilized embryos isolated from C57BL/6 mice. The resulting pups were genotyped by PCR with a mixture of three *Zbp1* primers (5’-CAACGACTGCTGCTGTCTTGC, 5’-GAACTCTGTGAAAGCCCTGTGAGG, 5’-CCTCATCCCCTGGTTGGTGTTAC). Mice with a WT genotype produce a 376 base pair band and the ZBP1^-/-^ genotype produce a 538 base pair band (Fig. S1B).

### Necroptosis assay

Lactate dehydrogenase (LDH) release was used as a measure of necroptosis in bone marrow derived macrophages (BMDM) from 2 independent experiments according to the manufacturer’s instructions (Sigma-Aldrich, MAK066-1KT). Bone marrow derived macrophages were harvested from 8 – 10-week-old mice and grown in 20% L929 conditioned RPMI medium as described (31). Harvested BMDM were seeded in technical triplicate at 5 x 10^4^ cells/well in a 96-well plate. Cells were infected with 2.5 x 10^5^ ME49 tachyzoites or left uninfected for 3 hours. After infection, the cells were washed with PBS (Calcium and magnesium free) and stimulated with 50 ng/mL TNF (PMC3013) and 40 μM Z-VAD-FMK (ab120382) or left unstimulated for 24 hours. Necrostatin-1 (SC-200142) was added at 50 μM as a control to inhibit necroptosis. Media supernatant was used to measure LDH release at 450 nm every 5 minutes for 1 hour in a BioTek Synergy HT plate reader. An NADH standard curve was used to determine the amount of LDH activity for each sample by the colorimetric assay.

### Parasites for in vitro and in vivo experiments

The ME49 type-2 *T. gondii* strain was used for all experiments and maintained in human foreskin fibroblast cells. The mCherry parasites were generated as follows: ME49 *T. gondii* strain (1 x 10^7^) was electroporated with 25 μg tub-mCherry (32), linearized with KpnI, and selected with chloramphenicol. Clones were isolated by limiting dilution. Tachyzoites were injected into mice for at least 28 days and bradyzoites were collected from the brains for passage through a feline and mCherry oocysts were collected from the feces. Oocysts concentration was calculated using a hemocytometer. Brain tissue cysts were collected from chronically infected C57BL/6J mice and the number of brain tissue cysts for oral infection was determined by immunofluorescence as described previously (24).

### Survival curve

Male or female mice between 9 and 13 weeks old were used for oral and intraperitoneal (IP) challenges. Oral challenges were performed by gavage with mCherry oocysts or bradyzoite brain cysts. The mCherry oocyst oral challenge was done with 6 x 10^3^ oocysts/mouse and the brain tissue cyst oral challenge was done with 2 x 10^3^ or 4 x 10^3^ brain tissue cysts/mouse. The IP infection was performed with 1 x 10^4^ tachyzoites/mouse. All mice were monitored daily for clinical signs of disease and euthanized when moribund.

### Intestine pathology

Intestine pathology was evaluated by histology and length. The Comparative Pathology Laboratory at UW-Madison scored blinded hematoxylin and eosin (H&E) stained ileum swiss rolls from female mice infected with 600 mCherry oocysts by oral gavage at day 7 post infection. The ileum (distal 1/3 of the small intestine) was washed with PBS and fixed in 10% buffered formalin in a swiss roll. Uninfected female mice ileum samples were processed as controls. The slides were blinded and scored from 0 – 5 (0 = equivalent to control and 5 = severe) on inflammation, gland loss, smooth muscle vacuolation, and edema. The intestine length was measured in male and female mice at 7 – 10 weeks old that were gavage fed 1 x 10^4^ mCherry oocysts. The small intestine was measured at 7 days post infection. Uninfected male and female mice 7 – 10 weeks old were used to normalize the intestine length of infected mice.

### In vivo parasite quantification

Parasite burden was determined in intestine samples by mCherry fluorescence and qPCR. The UW-Madison Small Animal Imaging and Radiotherapy Facility provided an *in vivo* imaging system (IVIS) to measure mCherry fluorescence. Male and female mice at 7 – 11 weeks old were gavage fed 1 x 10^4^ mCherry oocysts/mouse in 2 independent experiments. At 7 days post infection the mice were sacrificed, their intestines were removed, and mCherry fluorescence was measured (excitation 587 and emission 610) by IVIS. The fluorescence of infected WT and RIPK3^-/-^ mice was normalized to the average fluorescence in uninfected male and female mice. Parasite burden measured by qPCR was performed with parasite specific SAG1 primers (5’-TGCCCAGCGGGTACTACAAG and 5’-TGCCGTGTCGAGACTAGCAG) on gDNA from three intestine sections. Total gDNA was extracted from 1 cm sections from the duodenum, jejunum, and ileum tissues in female mice orally infected with 3 x 10^4^ mCherry oocysts at 7 days post infection by Trizol following the manufacturer’s instructions. A gDNA *T. gondii* standard curve was generated from a known concentration of parasites cultured in human foreskin fibroblast cells to calculate burden. A StepOne Real-Time PCR machine with iTaq Universal SYBR Green Supermix (Bio Rad, 1725120) was used to determine parasite number in each intestine section relative to a standard curve of parasite gDNA. Total parasite/ng was normalized across all 3 intestine tissue sections for each mouse sample.

### Brain cyst quantification

The number of brain cysts was determined by immunofluorescence at 28 days post oral infection with 3 x 10^3^ mCherry oocysts in male mice. Each brain was homogenized separately with a Dounce homogenizer, fixed with 4% paraformaldehyde for 20 minutes, quenched with 0.1 M glycine for 2 minutes, and blocked for 1 hour at room temperature in blocking buffer (PBS with 3% BSA and 0.2% Triton-X 100). Brain cysts were stained with streptavidin conjugated Dolichos biflorus agglutinin (Vector laboratories, B-1035-5) for 1 hour at room temperature, washed in PBS with 0.1% Triton-X 100, followed by incubation with a biotinylated Alexa Flour 594 (Thermo Fisher, S11227) secondary antibody for 1 hour at room temperature in the dark. An aliquot of processed brains was mounted on glass coverslips, blinded and the number of cysts was counted on a Zeiss inverted Axiovert 200 motorized microscope with 10x objective.

### In vitro parasite quantification

The parasite burden *in vitro* was determined by immunofluorescence in BMDM. A total of 1 x 10^5^ BMDM were seeded on glass coverslips and infected with 5 x 10^5^ tachyzoite parasites for 3 hours. After infection, cells were stimulated with 25 ng/mL LPS and 25 U/mL IFN-γ for 24 and 48 hours. Cells were fixed with 3% formaldehyde for 20 minutes, quenched for 5 minutes with 0.1M glycine, and blocked for 1 hour at room temperature in blocking buffer (PBS with 3% BSA and 0.2% Triton-X 100). Parasites were stained with chronic infection serum for 1 hour at room temperature, followed by Alexa Flour 488 anti-mouse (Thermo Fisher, Z25002) secondary antibody for 1 hour. All slides were blinded before counting. The 24-hour timepoint included 7 WT and 5 RIPK3^-/-^ coverslips, and the 48-hour timepoint included 6 WT and 6 RIPK3^-/-^ coverslips. Total parasites were counted in 150 vacuoles and visualized using a Zeiss inverted Axiovert 200 motorized microscope with a 100x oil objective.

### Intestine permeability assay

Intestine permeability was determined by LPS and FITC-dextran concentration in mouse blood serum after oral infection. Serum LPS concentration was measured with the Pierce Chromogenic Endotoxin Quant Kit (Thermo Scientific, A39552) following the manufacture’s protocol. Male and female mice were orally infected with 1 x 10^4^ mCherry oocysts by gavage. As a positive control, male and female mice were treated with 3% dextran sodium sulfate (DSS) in drinking water. Paired blood serum was collected at days 0 (uninfected), 3, 5, and 7 post infection. Blood serum LPS concentration was measured spectrophotometrically at 405 nm in a 96-well BioTek Synergy HT plate reader. Intestine permeability was determined by FITC-dextran concentration in mouse blood serum after oral infection. Female mice were infected for 7 days with 600 mCherry oocysts by gavage. Mice were fasted overnight, gavage fed 0.44 mg/g FITC-dextran, and after 4 hours blood serum was collected at euthanasia. FITC-dextran was quantified spectrophotometrically (485 nm excitation and 528 nm emission) from blood serum.

### Blood serum cytokine measurement

The BD Cytometric Bead Array Mouse Inflammation Kit (BD Biosciences, 552364) was used to measure IL-6, IL-10, monocyte chemoattractant protein-1 (MCP-1), IFN-γ, TNF, and IL-12 from blood serum. Female mice at 8 – 13 weeks old were gavage fed 6 x 10^3^ mCherry oocysts and euthanized at 7, 8, and 9 days post infection in 3 independent experiments. Blood serum was collected for cytokine analysis by flow cytometry with a ThermoFischer Attune at the UW-Madison Flow Cytometry Core.

## Results

### *ZBP1^-/-^* and *RIPK3^-/-^ mice show divergent phenotypes to necroptosis and host survival*

The ZBP1 knockout mice used in Pittman *et al*. 2016 (24) originated from the Shizuo Akira lab and appear to have a mixed genetic background (33). This mixed background in the Akira knockout has complicated comparisons to WT mice (33). To rectify this, we created a new ZBP1 null (ZBP1^-/-^) mouse in the C57BL/6 background using CRISPR-Cas9 technology to remove the entire *Zbp1* genomic locus, including the promoter and all mRNA splice variants (Fig. S1). We then evaluated the role of ZBP1 in host protection during *T. gondii* infection.

Necroptosis is a pro-inflammatory cell death pathway where cells become permeable and release damage associated molecular compounds that enhance inflammatory responses (20). Because ZBP1 has been shown to drive RIPK3-dependent necroptosis (34), we tested whether ZBP1 could activate this pathway during *T. gondii* infection. We included RIPK3 null (RIPK3^-/-^) mice as a negative control for necroptosis because RIPK3 kinase and scaffold domains are essential for necroptosis (35, 36). Necroptosis was measured in WT, ZBP1^-/-^, and RIPK3^-/-^ bone marrow derived macrophages (BMDM) by lactate dehydrogenase (LDH) release. LDH release was reduced in all genotypes treated with necrostatin-1, a necroptosis inhibitor (Fig. 1A). Infection with *T. gondii* alone caused little LDH release in each BMDM genetic background, but stimulation with TNF and Z-VAD-FMK (a pan caspase inhibitor that drives necroptosis) caused a similar increase in LDH release in WT and ZBP1^-/-^ BMDM. The LDH release in WT and ZBP1^-/-^ BMDM was amplified when stimulation with TNF and Z-VAD-FMK was combined with *T. gondii* infection (Fig. 1A). In contrast, RIPK3^-/-^BMDM showed no LDH release above background in any condition. This outcome indicates that ZBP1 does not promote cell permeability, a process apart of necroptosis, while RIPK3 influences cell membrane integrity during *T. gondii* infection.

**FIG 1.**
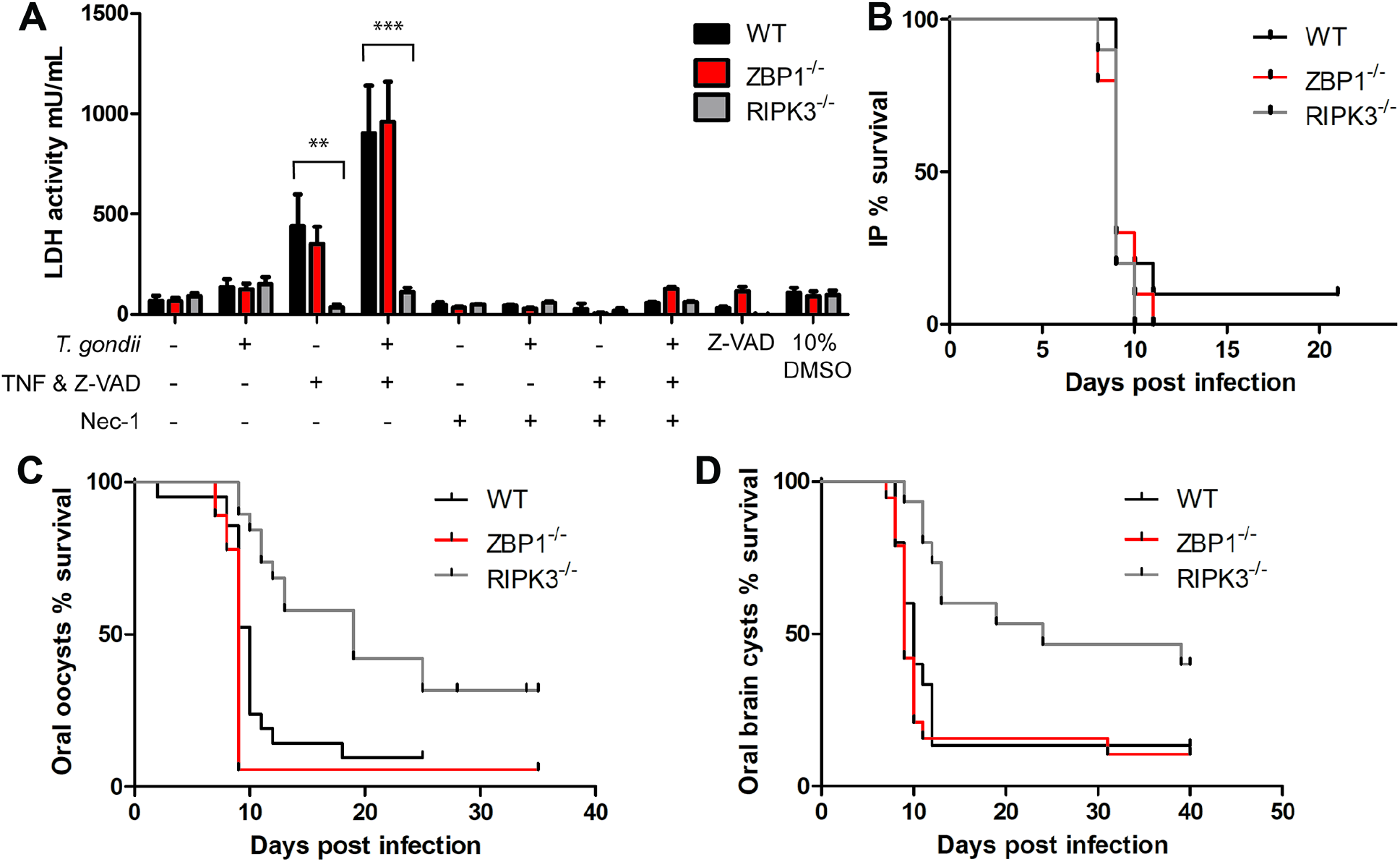
ZBP1 and RIPK3 show divergent phenotypes to necroptosis and host survival. (A) Necroptosis was measured by LDH release in BMDM. Cells were seeded at 5 x 10^4^ cells/well in triplicate in a 96-well plate and infected with 2.5 x 10^5^ parasites/well or left uninfected for 3 hours. Stimulated cells received 50 ng/mL TNF and 40 μM Z-VAD-FMK and unstimulated cells received fresh media. After 24 hours, the media supernatant was used to measure LDH release by absorption. Necrostatin-1 was added upon stimulation at 50 μM as a negative control for necroptosis. These results are from 2 independent experiments. A 2-way ANOVA with Bonferroni post-test was used for statistical analysis in LDH release. ** P-value < 0.01, *** P-value < 0.001. (B) Host survival from IP inoculation with *T. gondii* tachyzoites. Female (WT n = 10, ZBP1^-/-^ n = 10, RIPK3^-/-^ n = 10) mice were IP infected with 1 x 10^4^ tachyzoites/mouse. This data combines 2 independent experiments. Significance was determined by a Log-rank (Mantel-Cox) test and the P-value between each genotype was not significant. (C) Host survival from an oral infection with oocysts. Male (WT n = 17, ZBP1^-/-^ n = 12, RIPK3^-/-^ n = 15) and female (WT n = 9, ZBP1^-/-^ n = 6, RIPK3^-/-^ n = 8) mice were orally infected by gavage with 6 x 10^3^ mCherry oocysts/mouse in 3 independent experiments. Significance was determined by a Log-rank (Mantel-Cox) test. The P-value = 0.001 for WT compared to RIPK3^-/-^ mice and the P-value was not significant between WT and ZBP1^-/-^ mice. (D) Host survival from an oral infection with brain tissue cysts. Female (WT n = 15, ZBP1^-/-^ n = 17, RIPK3^-/-^ n = 15) mice were orally infected by gavage with 4 x 10^3^ brain tissue cysts in 3 independent experiments. Significance was determined by a Log-rank (Mantel-Cox) test. The P-value = 0.003 between WT and RIPK3^-/-^ mice and the P-value was not significant between WT and ZBP1^-/-^ mice.

Pro-inflammatory cell death pathways, like necroptosis, have evolved to protect the host from pathogenic infections (18). An acute host survival challenge is a simple assay to determine a gene’s contribution to host protection during infection. We established host protection against *T. gondii* by intraperitoneal (IP) and oral infection. Similar to previous observations (20), there was no significant difference between WT and ZBP1^-/-^ mouse survival after IP inoculation (Fig. 1B), but the ZBP1^-/-^ mice were more susceptible to oral *T. gondii* challenges with oocysts and brain tissue cysts (Fig. 1C&D, S2 - 4). Along with ZBP1^-/-^ mice, the RIPK3^-/-^ mice also showed no difference in survival after IP inoculation with *T. gondii* tachyzoites (Fig. 1B). However, unlike ZBP^-/-^ mice, the RIPK3^-/-^ mice had significantly improved survival after oral infection with both *T. gondii* oocysts and brain tissue cysts (Fig. 1C & D). These results suggest that while ZBP1 and RIPK3 both play roles during oral *T. gondii* infection, they are likely working in different pathways.

### RIPK3^-/-^ mice have higher parasite burdens

We sought to understand the role of RIPK3 in host defense responses given the striking survival phenotype to oral infection. Genetic evidence supports RIPK3-dependent necroptosis clears viral, bacterial, and parasitic infections which can affect host susceptibility (25, 26, 36–40). We determined the *T. gondii* parasite burden at the peak acute infection by measuring mCherry parasite fluorescence and by qPCR in intestine sections. A significantly higher mCherry signal was present in RIPK3^-/-^ intestines compared to WT (Fig. 2A) which correlated with a greater parasite load by qPCR (Fig. 2B). Oral infection with a sub-lethal mCherry oocyst dose also showed an increased parasite burden in RIPK3^-/-^ small intestine and livers during acute infection (Fig. S5) and mouse brains at chronic infection (Fig. 2C). Moreover, RIPK3^-/-^ BMDM exhibited increased total parasite burden by immunofluorescence *in vitro* after infection and stimulation (25 ng/mL LPS and 25 U/mL IFN-γ) (Fig. 2D). These results reinforce previous evidence for RIPK3 control of pathogenic infections, and more importantly, these results suggest that fatality in WT mice is not a consequence of increased parasite replication.

**FIG 2.**
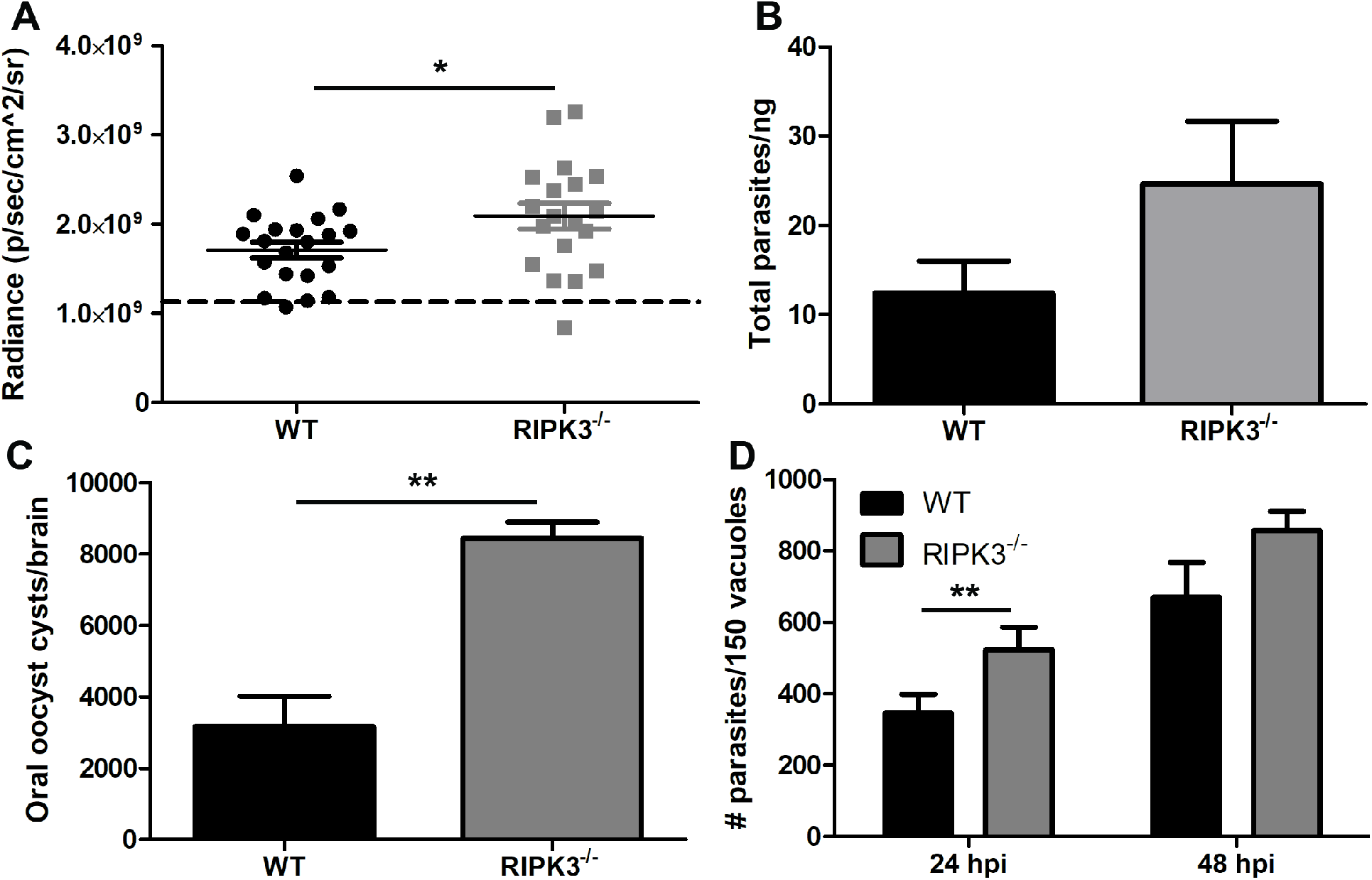
RIPK3^-/-^ mice have higher parasite burdens. (A) Intestine parasite burden determined by mCherry fluorescence with IVIS. Male (WT n = 10, RIPK3^-/-^ n = 9) and female (WT n = 10, RIPK3^-/-^ n= 10) mice were gavage fed 1 x 10^4^ mCherry oocysts and mCherry fluorescence was measured (excitation 587 and emission 610) in extracted intestines at 7 days post infection. The dotted line indicates the average background mCherry fluorescence of uninfected male (WT n = 2, RIPK3^-/-^ n = 2) and female (WT n = 4, RIPK3^-/-^ n = 3) mouse intestines. (B) Parasite burden measured by qPCR in intestines with parasite specific SAG1 primers. Total gDNA was extracted from 1 cm intestine sections (duodenum, jejunum, and ileum) in female (WT n = 3, RIPK3^-/-^ n = 3) mice orally infected with 3 x 10^4^ mCherry oocysts. A standard curve was generated from a known concentration of tachyzoite parasites to calculate the burden. The intestine parasite burden was normalized across all 3 intestine tissue sections for each mouse. P-value was not significant for the intestine qPCR. (C) Brain cyst burden in male (WT n = 5, RIPK3^-/-^ n = 3) mice 28 days post oral infection with 3 x 10 mCherry oocysts. (D) Parasite burden in WT and RIPK3^-/-^ BMDM. Total parasites in 150 vacuoles were counted by IFA on blinded glass coverslips at 24 (WT n = 7, RIPK3^-/-^ n = 5) and 48 (WT n = 6, RIPK3^-/-^ n = 6) hour post infection and stimulation (25 ng/mL LPS and 25 U/mL IFN-γ). A 2-tailed independent Student’s T-test was used to calculate significance. * P-value < 0.05, ** P-value < 0.01.

### WT and RIPK3^-/-^ mice do not have differences in intestinal villi pathology

As RIPK3 is associated with mucosal immune pathology (41, 42) and oral *T. gondii* infection causes intestinal inflammation (43–46), we assessed intestinal villi immunopathology. At day 7 post oral *T. gondii* infection, H&E stained ileum swiss rolls were blinded and assessed for villi integrity. While we saw signs of villi damage due to oral *T. gondii* infection, there were no differences in villi damage between WT and RIPK3^-/-^ mice (Fig. 3). We further examined villi integrity by measuring intestine permeability because it was previously seen that oral *T. gondii* infection causes lipopolysaccharide (LPS) from the bacterial microbiome as well as gavage fed FITC-dextran to enter the circulation (47). Using a dextran sulfate sodium-induced colitis model as a positive control, we measured LPS in the blood at 3, 5, and 7 days after oral *T. gondii* infection. There were no significant differences in serum LPS (Fig. 4A) or gavage fed FITC-dextran concentration in blood serum between WT and RIPK3^-/-^ mice (Fig. S6). These results suggest that intestine permeability induced by oral *T. gondii* infection is equivalent in both strains and the increased survival seen in RIPK3^-/-^ mice after oral *T. gondii* infection is not due to decrease intestinal villi pathology.

**FIG 3.**
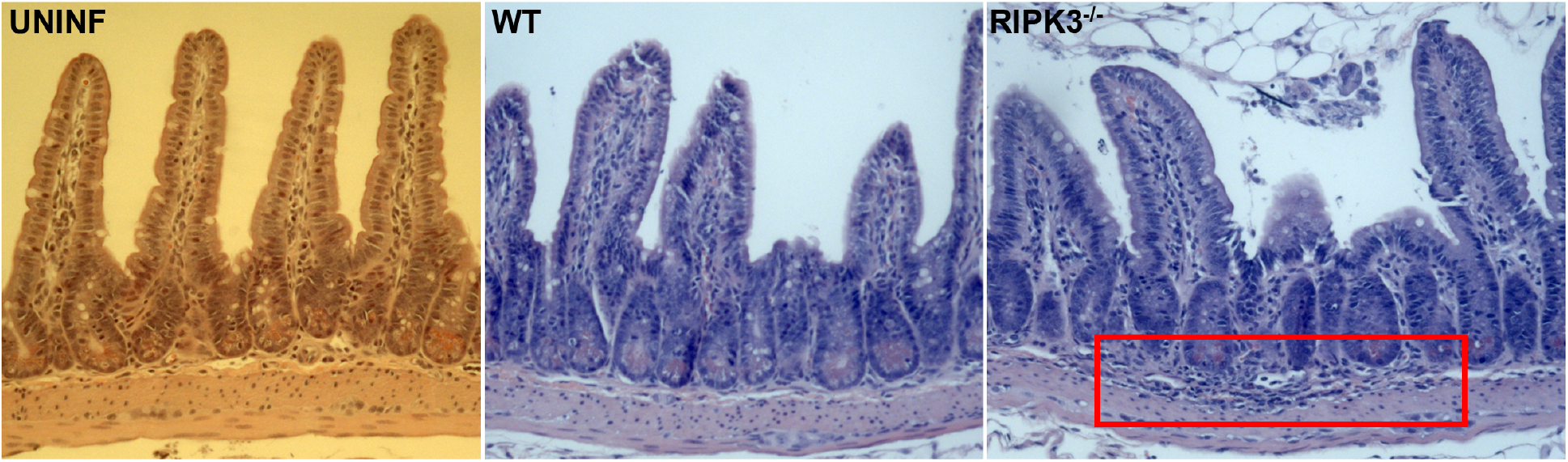
Intestinal histopathology shows similar villi damage in WT and RIPK3^-/-^ mice. Female mice were left uninfected (UNINF, left panel) or were infected with 600 mCherry oocysts by oral gavage and sacrificed at day 7 post infection (WT, middle panel and RIPK3^-/-^, right panel). The ileum (distal 1/3 of the small intestine) was washed with PBS and fixed in 10% buffered formalin in a swiss roll, sectioned, then stained with H&E and imaged using Zeiss light microscopy. The red box indicates dark nuclei of inflammatory cells that have infiltrated in the lamina propria and submucosa of this tissue.

**FIG 4.**
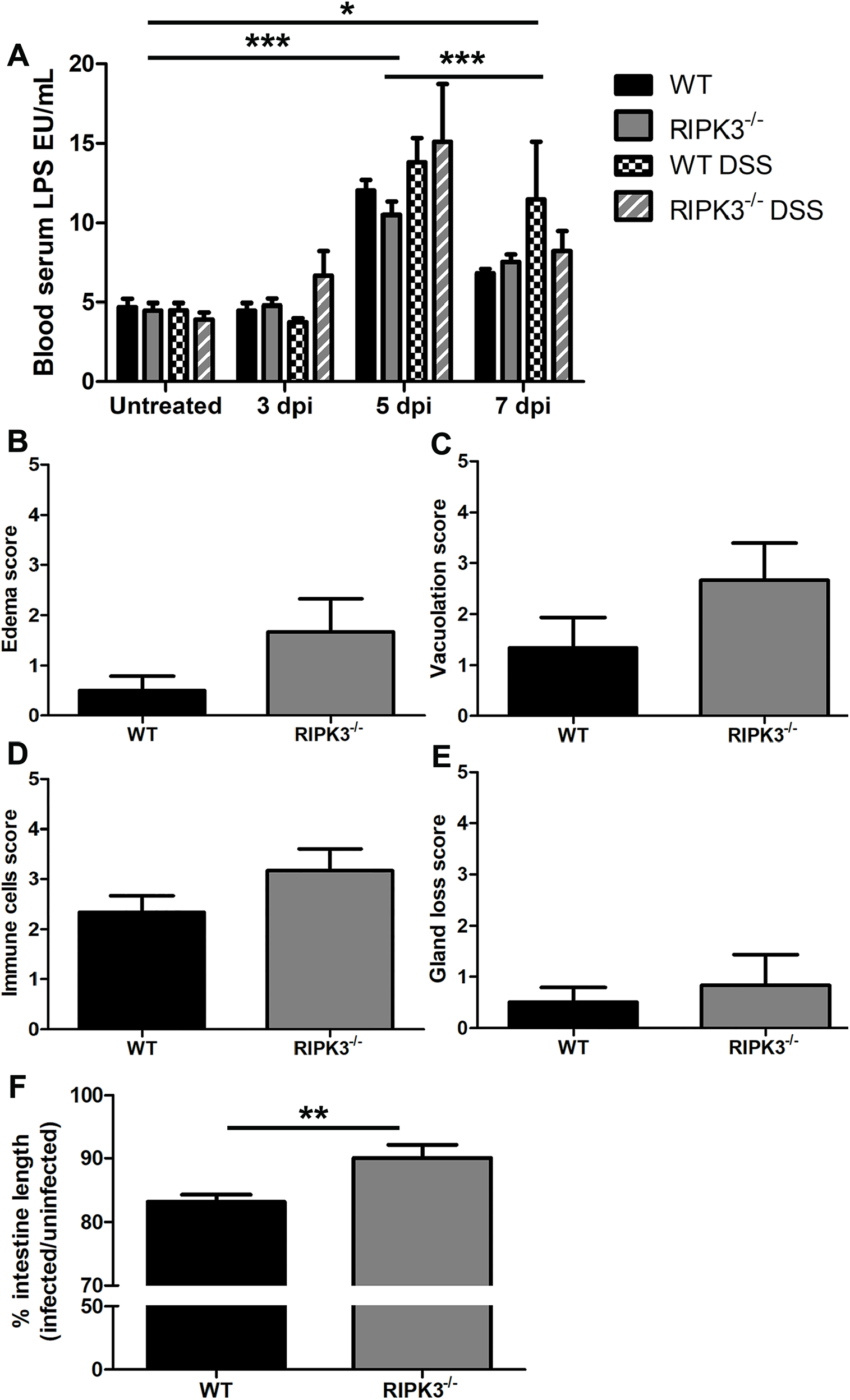
RIPK3 does not affect intestinal permeability but lamia propria pathology after oral T. gondii infection. (A) Intestine permeability by LPS concentration in blood serum. Male (WT n = 4, RIPK3^-/-^ n =3) and female (WT n = 5, RIPK3^-/-^ n = 5) mice were orally infected with 1 x 10^4^ mCherry oocysts by gavage. As a positive control, male (WT n = 4, RIPK3^-/-^ n = 5) and female (WT n = 4, RIPK3^-/-^ n = 3) mice were treated with 3% DSS in the drinking water. Paired blood serum was collected at days 0 (uninfected), 3, 5, and 7 post oral infection. Statistical significance was calculated by 2-way ANOVA with Bonferroni post-test. *** P-value < 0.001 for uninfected compared to 5 dpi, and 5 dpi compared to 7 dpi. * P-value < 0.05 for uninfected compared to 7 dpi. There was no significant difference within each group. (B-E) Intestine pathology scores by H&E in ileum swiss rolls. Female mice (WT n = 3, RIPK3^-/-^ n = 3) were infected with 600 mCherry oocysts by oral gavage and sacrificed at day 7 post infection. The ileum (distal 1/3 of the small intestine) was washed with PBS and fixed in 10% buffered formalin in a swiss roll. Uninfected female mice (WT n = 2, RIPK3^-/-^ n = 3) ileum samples were processed the same as controls. The slides were blinded and scored from 0 – 5 (0 = equivalent to control and 5 = severe) for (B) edema, (C) smooth muscle vacuolation, (D) immune cell infiltration, and (E) gland loss. (F) Intestine length was measured in male (WT n = 6, RIPK3^-/-^ n = 6) and female (WT n = 6, RIPK3^-/-^ n = 6) mice at 7 days post oral infection by gavage with 1 x 10^4^ mCherry oocysts. A 2-tailed independent Student’s T-test was used to calculate significance from 2 independent experiments. ** P-value < 0.01.

### RIPK3^-/-^ mice have more immune cell infiltration and edema in the lamina propria

After oral infection, *T. gondii* parasites can be found largely in the lamina propria of the ileum (48–50). We noticed that large patches of immune cells tended to be more common in the lamina propria of RIPK3^-/-^ mice after oral *T. gondii* infection (Fig. 3), so we submitted the H&E stained ileum swiss rolls to the UW-Madison Comparative Pathology Laboratory for analysis. They scored gland loss, immune cell infiltration, edema in the lamina propria, and vacuolation of the muscularis mucosa (Fig. 4B-E). While none of these scores reached statistical significance, there was a trend for the RIPK3^-/-^ mice to have increased inflammatory measures compared to WT mice. We then measured the length of the entire infected small intestine because intestine shortening is associated with tissue damage and pathology from oral *T. gondii* infections (43–46). The difference in intestine length was significant (Fig. 4F) as WT mice lost 17% intestine length compared to a 10% loss in RIPK3^-/-^ mice. The longer RIPK3^-/-^ intestine length could be due to swelling from increased edema, immune cell infiltration, and muscularis vacuolation or it could be due to decreased immunopathology.

### RIPK3^-/-^ mice have higher IFN-γ and lower IL-10

As inflammatory cytokines are critical for the control of *T. gondii* infection, we compared serum cytokine levels between WT and RIPK3^-/-^ mice using the cytometric bead array mouse inflammation kit. There were no differences in MCP-1, IL-6, IL-12, and TNF systemic cytokines levels between WT and RIPK3^-/-^ mice at any of the time points examined (Fig. S7A-D). However, at day 9 post oral infection, RIPK3^-/-^ mice had significantly higher IFN-γ (Fig. 5A) and reduced IL-10 levels (Fig. 5B). IFN-γ and IL-10 are both essential for mice to survive acute *T. gondii* infection, but IFN-γ is necessary to control parasitemia (9, 10) and IL-10 is necessary to control the host immune response (13, 14). These results may be due to the higher parasitemia in the RIPK3^-/-^ mice (Fig. 2).

**FIG 5.**
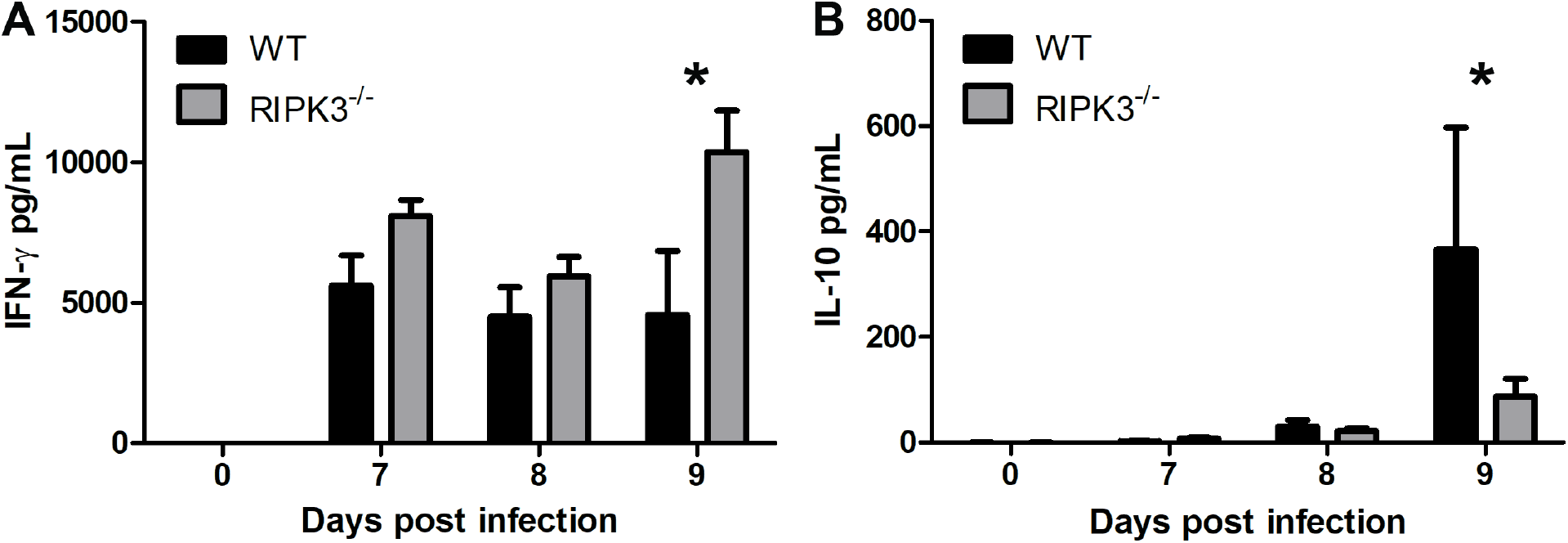
*RIPK3 activity* affects IFN-γ and IL-10 serum cytokine levels. Blood serum cytokines in female (WT n = 14, RIPK3^-/-^ n = 15) mice gavage fed 6 x 10 mCherry oocysts. Serum samples were collected at 7, 8, and 9 days post infection in 3 independent experiments. IFN-γ (A) and IL-10 (B) were significantly different. A 2-way ANOVA with Bonferroni post-test was used to calculate significance. * P-value < 0.05.

### *MLKL*^-/-^ and *RIPK3^-/-^ mice show divergent host survival phenotypes*

Necroptotic induced cell death and inflammation are dependent on RIPK3, but RIPK3 activity is not limited to necroptosis (51). The necroptosis executioner, MLKL, is the downstream substrate of RIPK3 that ultimately compromises cell membranes, fulfilling the necroptotic pathway (52). We specifically examined RIPK3-dependent necroptosis in host survival to oral *T. gondii* infection with MLKL null (MLKL^-/-^) mice. After oral infection, MLKL^-/-^ mice succumb to infection similar to WT mice, whereas RIPK3^-/-^ mice showed improved survival (Fig. 6, S8). This data provides evidence that RIPK3-independent necroptosis contributes to severe intestine pathology and host death.

**FIG 6.**
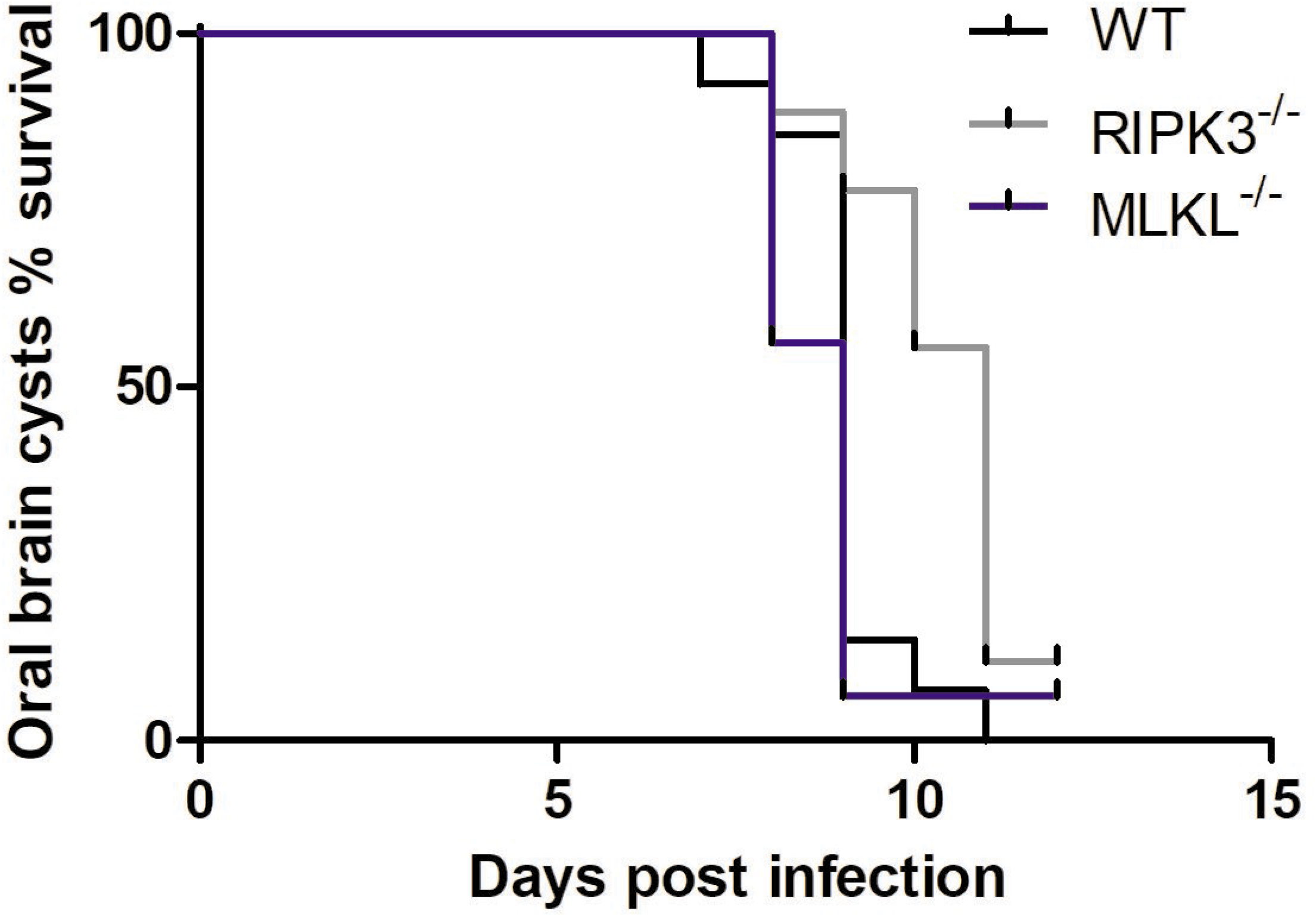
RIPK3-independent necroptosis activity influences host survival to oral. Female (WT n = 25, RIPK3^-/-^ n = 16, MLKL^-/-^ n = 16) mice were orally infected with 4 x 10^3^ brain tissue cysts by gavage. A Log-rank (Mantel-Cox) test was used to calculate significance between each genotype. ** P-value <0.01. The results are a compilation of 3 independent experiments.

## Discussion

Innate immune activation and its role in host defense to pathogenic infections depend on PRR driven inflammation. ZBP1 has recently come to light as a PRR involved in innate immunity and programmed cell death (34). Mechanisms of programmed cell death include apoptosis, necroptosis, and pyroptosis. *T. gondii* has evolved mechanisms to inhibit apoptosis to promote fitness (53). However, the host maintains cellular cross-talk between each programmed cell death pathway to subvert apoptosis inhibition and guard against pathogenic infection via necroptosis and pyroptosis (18). ZBP1 was found to mediate the interferon-induced necroptosis pathway in response to viral infection (27, 28). ZBP1 as a PRR that induces immunity could fit within the realm of bacterial and parasitic infection (24, 54–56). The fact that IFN-γ and ZBP1 are highly expressed and maintained throughout *T. gondii* infection (22, 23), make *T. gondii* a thought-provoking target to investigate the role of ZBP1 in non-viral infections. Our lab previously found ZBP1 to be important for oral but not IP infection (24). Here, we sought to determine the potential role of ZBP1 in host induced necroptosis in response to *T. gondii* infection by comparing ZBP1^-/-^ and RIPK3^-/-^ mice.

We tested ZBP1 induced necroptosis by LDH release in BMDM and found ZBP1 to be independent of necroptosis in *T. gondii* infection. We also found ZBP1^-/-^ and RIPK3^-/-^ mice to have opposing responses to oral *T. gondii* infection, with ZBP1^-/-^ mice being more susceptible (24) and RIPK3^-/-^ mice being more resistant (Fig. 1). Studies using high dose oral *T. gondii* infection have linked excessive Th1 immunity and intestinal pro-inflammatory cell death as the culprit in host mortality (15–17). The physiological role of RIPK3-dependent necroptosis can drive immunopathology and impair host fitness (35, 36, 41, 42). These findings support the fact that RIPK3^-/-^ mice have improved survival to oral *T. gondii* infection even with a higher parasite burden (Fig. 2). While there were no differences between WT and RIPK3^-/-^ mice in intestinal villi damage, there was a trend for the RIPK3^-/-^ mice to have increased inflammatory measures in the lamina propria (Fig. 3, 4), as well as a reduction in IL-10 (Fig. 5). IL-10 is up-regulated to control excessive Th1 cells and prevent immunopathology (57) and has previously been seen to be key in host survival after *T. gondii* infection (13, 14). All of these factors contradict a model where RIPK3^-/-^ mice survive oral *T. gondii* infection better than WT mice because of reduced immunopathology. RIPK3^-/-^ mice have reduced intestinal length loss after oral *T. gondii* infection compared to WT mice (Fig. 4F). This difference could be due to the WT mice having RIPK3-dependent activity causing more severe intestinal immunopathology, or more likely, it could be that the small intestine in the RIPK3^-/-^ mice is more swollen and longer with the increased edema, immune cell infiltration, and muscularis vacuolation (Fig. 4B-D).

Gut bacteria drive an unregulated immune response that results in early host death, cellular necrosis, and severe tissue pathology in susceptible C57BL/6 WT mice in high dose oral *T. gondii* infections (44, 58). The inflammatory response reduces antimicrobial compound secretion and compromises the epithelial barrier, which allows greater interaction with gut bacteria to promote Th1 immunity and pathogen control (47). Gnotobiotic and antibiotic treated mice are one of the few models that have also shown improved survival and higher parasite burdens after lethal oral *T. gondii* infections (44), similar to our RIPK3^-/-^ mice. Therefore, we examined intestine permeability by measuring bacterial LPS or gavage-fed FITC-dextran concentration in blood serum after oral *T. gondii* infection. There was no difference in LPS or FITC-dextran concentration in blood serum between WT and RIPK3^-/-^ mice (Fig. 4A, S6), which suggests intestine permeability and the degree of immune activation by gut bacteria is equivalent.

The biological function of RIPK3 appears to be only important for the natural route of *T. gondii* infection as there is not a difference in susceptibility between WT and RIPK3^-/-^ mice after IP infections. This implicates the importance of exposure route when studying host responses to pathogenic infections. The difference in RIPK3^-/-^ mouse survival between IP and oral infection likely contrast in pathogen recognition pathways that lead to immune stimulation. This result was evident in TLR-11 null (TLR-11^-/-^) mice that also have improved survival to oral but not IP *T. gondii* infection (59). TLR-11 in mice is a PRR that recognizes *T. gondii* profilin which activates MyD88 dependent signaling pathways to induce IL-12 and IFN-γ, two critical cytokines for host survival to *T. gondii* infection (60). When TLR-11^-/-^ mice are challenged with *T. gondii* by IP infection, the host is unable to recognize the pathogen, mount an appropriate immune response, and as a consequence, is overcome by uncontrolled parasite replication. In contrast, oral infection generates a TLR-11-independent protective immune response from gut bacteria that improves host survival. This study points towards an important role for gut bacteria to stimulate PRRs and immunity in the intestine to help fight pathogenic infections.

The role of RIPK3 is not limited to necroptosis. RIPK3 can activate pyroptosis as an alternative cell death mechanism that releases the pro-inflammatory effectors, IL-1β and IL-18. Pyroptosis and its pro-inflammatory effectors are activated upon *T. gondii* infection and have been shown to control parasite burden in *in vitro* and *in vivo* oral infection models (61, 62). The proportion of RIPK3 activity in necroptosis or pyroptosis in host survival to oral *T. gondii* infection was clarified with MLKL^-/-^ mice. The observation that MLKL^-/-^ mice succumb to oral infection with WT mice indicated that necroptosis does not play a major role in the survival difference between WT and RIPK3^-/-^ mice. The difference in survival between RIPK3^-/-^ and MLKL^-/-^ mice could be attributed to their ability to initiate pyroptosis because RIPK3 can activate pyroptosis in an MLKL-dependent or -independent manner (66, 67). This result agrees with previous findings that show inhibition of pyroptosis effectors IL-1β and IL-18 improve host survival to oral *T. gondii* infection (17, 65, 68, 69). However, genetic knockouts in other inflammasome and pyroptosis components have shown conflicting results in host susceptibility from IP *T. gondii* infection (70–72). Our study provides further evidence that supports inflammasome activation in oral *T. gondii* infection affects the host immune response.

Our CRISPR-Cas9 ZBP1^-/-^ mouse had no significant effect on host survival to IP infection and reduced survival to oral challenge. This result corresponded with our previous findings, where the Akira ZBP1 knockout mice also showed no significant difference to IP infection but reduced survival to oral infection (24). Although independent groups have linked ZBP1 function to necroptosis, they have reported opposing susceptibility phenotypes to viral infection with the Akira ZBP1 knockouts (26, 37). Thus, the relevance of ZBP1 to host vulnerability in infectious diseases could be remedied by our clean CRISPR-Cas9 ZBP1^-/-^ mouse.

## Acknowledgments

We sincerely thank JP Dubey for oocyst production of mCherry expressing parasites, Sarah Wilson for assistance with IVIS, Laura Lettenberg for assistance with parasite replication assays, Wynne Moss for the creation of the ME49 strain expressing mCherry, and Melissa Graham for blinded histopathological analysis. We also thank Genentech for providing RIPK3^-/-^ mice and Dr. Doug Green at St. Jude Children’s Research Hospital for providing (MLKL^-/-^) mice. This work was supported by the National Institutes of Health grants T32 AI007414 (PWC) and R01AI144016-01(LJK) as well as the National Science Foundation Graduate Research Fellowship Program grant DGE-1747503 (PWC). Any opinions, findings, and conclusions or recommendations expressed in this material are those of the authors and do not necessarily reflect the views of the funding agencies.

